# Generative and discriminative recurrence employ opposing strategies for robust vision

**DOI:** 10.64898/2026.07.02.736066

**Authors:** Lea-Maria Schmitt, Maartje Koot, Micha Heilbron, Floris P. de Lange

## Abstract

Recurrence is thought to enhance the robustness of biological vision, but how it achieves this feat is largely unknown. Perceptual robustness can be implemented through either lateral connections supporting local integration within a processing stage or feedback connections drawing on broader context from higher stages, and through either a discriminative objective optimising task-relevant classification or a generative objective learning to reconstruct the causes of visual input. But do these different types of recurrence engage distinct computational strategies? As this question is difficult to test in vivo, we endowed convolutional neural networks with varying recurrent architectures and training objectives, and evaluated the consequences for internal representations and behaviour across noise levels. Two distinct computational strategies emerged. Generative feedback followed a reductionist strategy, with representations becoming lower-dimensional through denoising, achieving robustness at moderate noise levels without noise training. Both discriminative lateral and feedback recurrence followed an expansionist strategy, increasing dimensionality to sharpen discriminability without denoising, but requiring noise training to achieve robustness. These dissociable signatures reflect fundamentally different computational mechanisms of robust vision and provide testable predictions for which form of recurrence the brain employs.

## 1 Introduction

The perception of the world around us is commonly challenged by external noise, as when raindrops on a window obscure the view of whatever lies behind. Despite this challenge, we may take a split second longer to recognise what is there [Tang et al., 2014], but will rarely fail. A prevailing computational account of this robustness of vision attributes it to a two-stage process in the ventral visual pathway of the brain: a feedforward pass that rapidly maps sensory input to an initial best guess [VanRullen, 2007], followed by iterative refinement of this interpretation through recurrent processing [van Bergen and Kriegeskorte, 2020]. Yet the nature of this recurrence — both its type and objective — remains unresolved.

Addressing this question with neuroscientific methods alone is notoriously difficult, as recurrent connections are so deeply intertwined that they resist selective manipulation, and it is unclear what measurable neural signature the objectives they implement would leave. Artificial neural networks Preprint. (ANNs) offer a tractable solution [Doerig et al., 2023, de Lange et al., 2022], which allows the type of recurrent architecture and training objective to be manipulated in silico, with their consequences for neural representations serving as testable predictions for the brain.

In terms of neural architecture, one candidate connection is lateral recurrence, which is thought to disambiguate noisy representations within a brain region (or layer in an ANN) by locally integrating activity across neighbouring neurons [Lamme and Roelfsema, 2000, Spoerer et al., 2017]. In support of its functional relevance, ANNs extended with lateral recurrence have been shown to improve alignment with brain responses [Kar et al., 2019, Rajaei et al., 2019] and human behavioural performance [Tang et al., 2018, Seijdel et al., 2021] under challenging conditions. Another candidate is feedback recurrence, which is thought to disambiguate noisy representations in lower brain regions by constraining their interpretation with more global information from higher brain regions [Lee and Mumford, 2003, De Lange et al., 2018]. Although feedback is known to support robust recognition in the brain as well [Kar and DiCarlo, 2021], it has not been directly pitted against lateral recurrence (but see Lindsay et al. [2022]). Do the fundamentally different mechanisms of lateral and feedback recurrence translate into distinct consequences for neural representations and behaviour under noisy viewing conditions?

Beyond the type of recurrent architecture, another distinction is the objective of recurrence, the task(s) it is trained to perform. The brain may follow different objectives that guide visual perception [Peters et al., 2024]. One candidate is a discriminative objective that tunes representations to best support decisions relevant for behaviour such as object classification [DiCarlo et al., 2012, Yamins and DiCarlo, 2016]. Under this objective, recurrence provides additional computational depth that can be deployed adaptively. Another candidate is a generative objective, which recruits recurrence to reconstruct input at lower regions from the prior world knowledge of higher regions [Rao and Ballard, 1999, Friston, 2005]. Both objectives have been independently shown to capture signatures of visual processing in the brain [Csikor et al., 2025, Kubilius et al., 2019], but how do the two directly compare in shaping representations for robust vision?

Here we systematically compared, as a function of noise in natural images, behavioural performance and neural representations (see Section 2 for a motivation of metrics) between ANNs varying in their type of recurrent connection and training objective. Our main contributions are:

- demonstrating reductionist coding in generative feedback, with denoising and dimensionality reduction under noise, and expansionist coding in discriminative recurrence
- demonstrating that generative feedback achieves noise robustness without noise training, whereas discriminative recurrence requires exposure to noisy input
- tracing these differences to dissociable representational signatures, which can be used to guide experiments on recurrence in the brain

## 2 Related Work

A longstanding idea in both computer and biological vision research is that perception involves generative inference: explaining sensory input by reconstructing it from an internal model of the world [Grenander, 1993, Yuille and Kersten, 2006, Peters et al., 2024]. In neural terms, this is often mapped onto feedback recurrence, in which higher regions predict or reconstruct activity in the regions below [Lee and Mumford, 2003, Rao and Ballard, 1999, Bastos et al., 2012]. Critically, because generative objectives are meant to enable capturing the causal structure of the environment, they are thought to yield representations that generalise across tasks and afford superior robustness compared to discriminative objectives tailored to narrow task-relevant decisions [Yuille and Kersten, 2006, Lee and Mumford, 2003].

In contemporary computer vision, generative ANNs have become increasingly powerful, for example, in diffusion-based image generation [Ho et al., 2020, Rombach et al., 2022] and as a pretraining objective [He et al., 2022]. Accumulating evidence suggests that such generative ANNs exhibit striking robustness [Jaini et al., 2024, Wiedemer et al., 2025, Gabeur et al., 2026], consistent with the view that generative objectives yield more transferable representations than discriminative alternatives. However, the architectures of these state-of-the-art generative approaches diverge too far from the hierarchical, recurrent, and local nature of visual cortex to serve as models of biological vision.

A more brain-inspired approach equips feedforward convolutional neural networks (CNNs) with recurrent connections and trains them with either a discriminative [Kubilius et al., 2018, Spoerer et al., 2017] or generative objective. For the latter, we adopted Predify, a framework grounded in the influential neuroscientific theory of predictive coding [Bastos et al., 2012], which augments CNNs with generative feedback connections [Choksi et al., 2021]. Choksi et al. [2020] demonstrated that this augmentation promotes more robust vision (see Fang et al. [2022] for an auditory analogue), although without comparison to other types of recurrent architectures and training objectives.

This comparison was undertaken by Lindsay et al. [2022], who found that discriminative networks with lateral or feedback recurrence outperformed a generative network on noisy inputs, while generative feedback exhibited stronger denoising and greater dimensionality reduction. However, those networks were evaluated on MNIST, where limited complexity may hamper generalisation. Moreover, discriminative networks were trained on noisy images, whereas the generative network was not, conflating objective with training regime. Here, we go beyond these limitations by investigating natural images spanning 100 categories and systematically varying both objective and training regime to provide a controlled assessment of their contributions to robust vision.

## 3 Methods

All code and data necessary to replicate the results will be made publicly available at https://doi.org/10.34973/5j08-5356 upon publication.

### 3.1 Networks

ANNs were built, trained, and evaluated in Python (version 3.10) with PyTorch [Paszke et al., 2019, version 1.13.1] on an NVIDIA GPU, with a minimum storage requirement of 35 GB. We extended a feedforward backbone trained on natural images with varying recurrent architectures (feedback vs lateral) and training objectives (discriminative vs generative): a generative feedback network, a discriminative feedback network, and a discriminative lateral network (Figure 1A).

**Figure 1:**
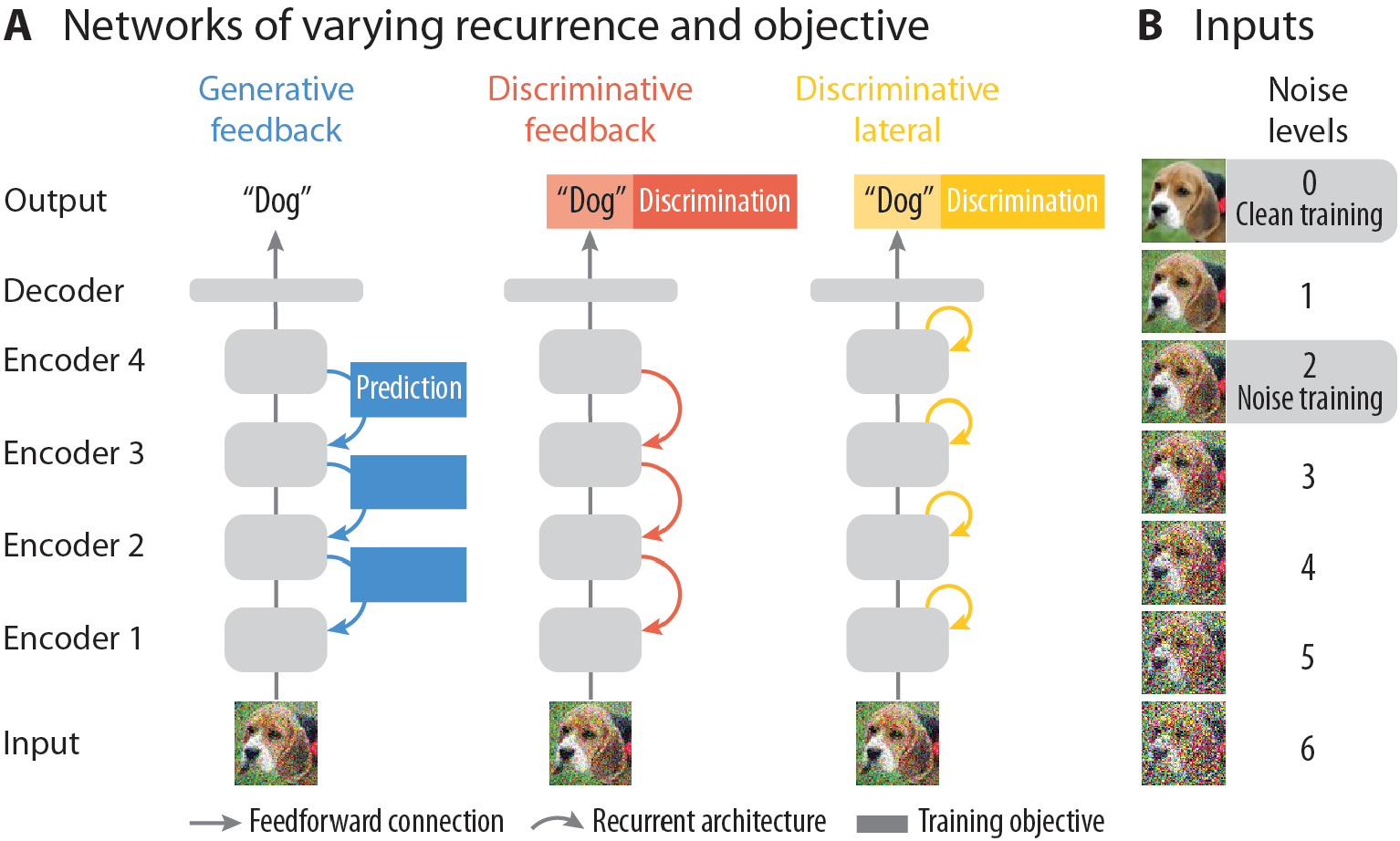
Networks and inputs. (A) A feedforward network with four convolutional encoders was first trained to classify clean images, then extended with varying recurrent architectures and training objectives: feedback connections trained to predict activations at the encoder below (generative feedback, blue), feedback connections trained to classify objects (discriminative feedback, red), or lateral connections trained to classify objects (discriminative lateral, yellow). (B) Network extensions were trained on either clean (noise level 0) or moderately noisy natural images (noise level 2) and tested across six levels of increasing pixel noise.

#### 3.1.1 Inputs

We trained and tested ANNs on MiniEcoset [Thorat et al., 2023], a subset of ecoset [Mehrer et al., 2021, CC BY-NC-SA 2.0]. The dataset consists of 235,000 natural images spanning 100 object categories, thereby striking a balance between dataset diversity and training efficiency. Images are 224 × 224 pixels with three colour channels (RGB).

Additionally, to probe recognition under challenging viewing conditions, we created noisy versions of images by applying Gaussian pixel noise to each pixel independently. For every image, we obtained six versions of increasing noise intensity by sampling pixel noise from Gaussian distributions with standard deviations *σ*∈{ 0.5, 1.0, 1.5, 2.0, 2.5, 3.0} (Figure 1B). To distribute noise equally across colour channels, the per-channel standard deviation was set to the total noise standard deviation divided by the square root of the number of channels.

#### 3.1.2 Feedforward backbone

The feedforward backbone followed the architecture of CORnet-Z [Kubilius et al., 2018], a shallow CNN designed to model processing along the ventral visual pathway during core object recognition. The network propagated the input through four stacked convolutional encoders in a strictly feedforward manner from lower to higher encoders. Encoders convolved their input using 64, 128, 256, and 512 filters respectively (encoder 1: kernel 7 × 7, stride 2, padding 3; encoders 2–4: kernel 3 × 3, stride 1, padding 1), yielding activations **a**^*e*^ at encoder *e*:

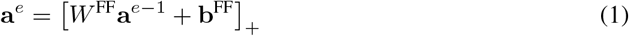

where *W* ^FF^ are convolutional feedforward weights, **b**^FF^ is the feedforward bias, and [·]_+_ is the ReLU nonlinearity. Max pooling (kernel 3 × 3, stride 2, padding 1) was applied after ReLU, such that **a**^*e*−1^ refers to the post-pooling activation of the preceding encoder.

In the decoder, activations from the final encoder were collapsed to a 512-dimensional vector via global average pooling, then linearly projected to 100 object categories. The network was trained to minimise cross-entropy loss between predicted and true class labels, constituting a purely discriminative objective. Following training, all feedforward weights and biases were frozen for subsequent training of recurrent network extensions.

#### 3.1.3 Generative feedback network

The discriminative feedforward backbone was extended with generative feedback following the Predify (version 0.0.1, GPL-3.0) implementation of Choksi et al. [2021]. Feedback connections were added from each encoder to the encoder directly below and trained to reconstruct lower-encoder activations from higher-encoder activations, constituting a generative objective. After an initial feedforward pass, activations evolved over time steps *T* according to:

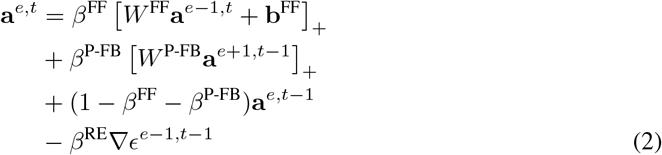

where the four terms correspond to: (i) the feedforward drive from the encoder below at the current time step; (ii) feedback from the encoder above at the previous time step, with feedback weights *W* ^P-FB^; (iii) a memory term retaining the encoder’s own activations from the previous time step; and (iv) a reconstruction error term where 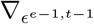 is the gradient of the MSE between the feedback prediction and the actual activations at the previous time step and encoder, correcting the current encoder’s representation toward generating more accurate predictions of the encoder below. The terms were balanced by coefficients *β*^FF^, *β*^P-FB^, and *β*^RE^. The frozen decoder continued to produce class labels, such that classification performance reflected the quality of feedback-modified encoder activations.

#### 3.1.4 Discriminative feedback network

As a further feedback variant, the feedforward backbone was extended with discriminative feedback, incorporating top-down connections from each encoder to the encoder directly below. Unlike the generative feedback network, this extension was trained on the same classification objective as the feedforward backbone. After an initial feedforward pass, activations evolved according to:

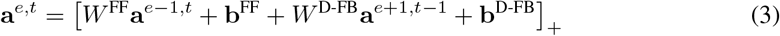

where *W* ^D-FB^ are feedback weights and **b**^D-FB^ is the feedback bias.

#### 3.1.5 Discriminative lateral network

As another network variant, the feedforward backbone was extended with lateral recurrence, incorporating self-connections at each encoder that refine activations from the previous time step. This extension was trained on the same classification objective as the feedforward backbone. After an initial feedforward pass, activations evolved according to:

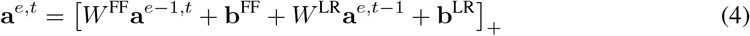

where *W* ^LR^ are lateral recurrence weights and **b**^LR^ is the lateral recurrence bias.

#### 3.1.6 Training

The feedforward backbone was trained separately from four different random seeds, once on clean images and once on images at noise level 2. All four clean-trained backbone instances served as the base for each network extension, which were likewise trained once on clean images and once on images at noise level 2. All network extensions were run for 13 time steps, comprising an initial feedforward pass followed by 12 recurrent time steps. Weights remained constant over time steps.

All networks were trained on the MiniEcoset training set. The feedforward backbone was trained in epochs of 917 batches of 256 images each, and the network extensions in epochs of 7,343 batches of 32 images each. The order of images was shuffled across epochs, and pixel noise was resampled for every presentation of each image. Weights and biases were initialised with He (Kaiming uniform) and updated using Adam [Kingma and Ba, 2014] with L2 regularisation. The learning rate and L2 penalty of all networks as well as the balancing coefficients of the generative feedback network were optimised using Optuna (v3.6.1, MIT License, Akiba et al., 2019; see Table 1 for a summary of hyperparameters). Training continued until no improvement in validation loss was observed over a minimum of 10 epochs.

**Table 1:** Training hyperparameters for all networks. Balancing coefficients *β*^FF^, *β*^P-FB^, and *β*^RE^ are reported per encoder. *β*^P-FB^ is 0 at encoder 4 as there is no higher encoder to receive feedback from, and *β*^RE^ is 0 at encoder 1 as there is no lower encoder to predict. Dashes indicate parameters not applicable to that network.

| Network | Learning rate | L2 penalty | $\beta^{\text{FF}}, \beta^{\text{P-FB}}, \beta^{\text{RE}}$ |
| --- | --- | --- | --- |
| Feedforward backbone | $1.90 \times 10^{-4}$ | $4.55 \times 10^{-3}$ | — |
| Generative feedback | $4.47 \times 10^{-5}$ | $4.23 \times 10^{-5}$ | $\begin{pmatrix} 0.37 & 0.12 & 0.00 \\ 0.96 & 0.01 & 0.61 \\ 0.89 & 0.02 & 0.04 \\ 0.23 & 0.00 & 0.11 \end{pmatrix}$ |
| Discriminative feedback | $1.66 \times 10^{-4}$ | $1.50 \times 10^{-3}$ | — |
| Discriminative lateral | $1.56 \times 10^{-5}$ | $3.65 \times 10^{-2}$ | — |

### 3.2 Analyses

All networks were evaluated on the held-out test set of 5,000 clean images and their noisy versions. As the feedforward backbone performs a single forward pass, it was evaluated after this pass. Recurrent network extensions were evaluated at the time step yielding the greatest gain in classification accuracy relative to the feedforward backbone, averaged across noise levels with the best-performing noise level weighted eight times (generative feedback: *t* = 12; discriminative feedback: *t* = 2; lateral recurrence: *t* = 4 (clean-trained) and *t* = 2 (noise-trained)).

Behavioural performance (top-1 classification accuracy) and neural representations (denoising and dimensionality) were quantified separately for each network at each noise level. The clean-trained feedforward backbone served as a baseline against which clean-trained network extensions were compared, isolating the unique contribution of each combination of recurrent architecture and training objective. The same analyses were subsequently carried out on noise-trained network extensions to test for any interactions between networks and training data.

#### Denoising

Network activations were extracted following an encoder’s nonlinearity. Representational denoising was quantified by calculating the Pearson correlation between encoder activations for each noisy image and its clean counterpart, with higher correlation indicating greater similarity between clean and noisy representations. A network extension was considered to have denoised its representations when its correlation exceeded that of the feedforward baseline at the same noise level.

#### Dimensionality

Representational dimensionality was quantified by performing principal component analysis across all image activations of an encoder, and then computing the area under the cumulative explained variance curve across all 5,000 principal components. A smaller area under the curve indicates that variance is more evenly distributed across components, reflecting higher-dimensional representations.

## 4 Results

We extended a feedforward backbone with different types of recurrent architectures and training objectives to investigate their behavioural and representational consequences for object recognition in noisy conditions (Figure 1).

### 4.1 Discriminative recurrence improves classification at low noise, generative feedback improves it at intermediate noise

We first established a feedforward baseline by training a network with four convolutional encoders to classify clean natural images. The network reached a top-1 classification accuracy of 0.47 (chance level: 0.01), in line with comparable networks trained on the same data [Thorat et al., 2023]. When tested on images degraded by increasing pixel noise, classification accuracy declined monotonically, with the steepest drop at intermediate noise levels and near-chance performance at the highest noise level (Figure 2, left), even when the network was trained on noisy images (Figure 4, left). This replicates the well-documented vulnerability of feedforward CNNs to noise in the input signal [Geirhos et al., 2018].

**Figure 2:**
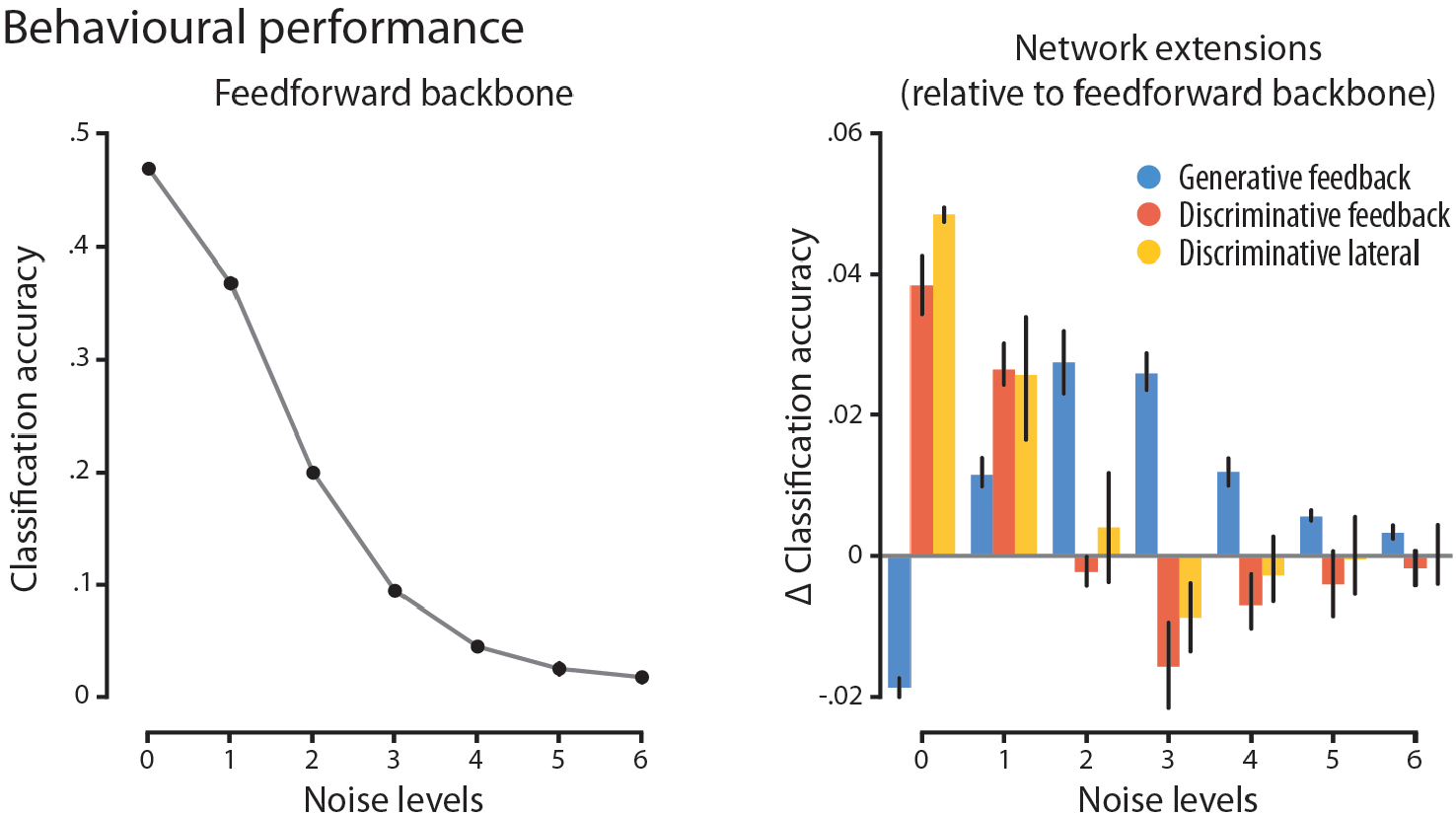
Behavioural performance. Top-1 classification accuracy of the feedforward backbone across increasing levels of pixel noise (left). Classification accuracy of recurrent extensions relative to the feedforward baseline (right). Error bars represent 95% confidence interval. Discriminative recurrence gains performance at low noise levels, generative feedback at intermediate noise levels.

Mirroring the delayed emergence of recurrence in brain development [Burkhalter, 1993], we then added either lateral or feedback connections trained with either a discriminative or generative objective on top of the frozen feedforward backbone. This allowed us to isolate the specific contributions of recurrent architecture and training objective over and above feedforward processing. Both discriminative networks improved classification accuracy for noise-free and mildly degraded images compared to the feedforward baseline, but performed worse for moderately degraded images. In contrast, the generative feedback network outperformed the feedforward baseline for moderately degraded images, while performing worse when images were either noise-free or severely degraded (Figure 2, right). Together, these results reveal that the training objective determines the degree of image degradation at which recurrence benefits recognition most.

### 4.2 Generative feedback but not discriminative recurrence denoises representations of noisy images

Having established how the type of recurrent architecture and training objective affects behavioural performance, we next asked how it shapes the underlying neural representations. We extracted activations from the first encoder, which receives only the image as feedforward input and is therefore free of upstream recurrent influences, to probe how recurrence reshapes feedforward representations directly. We then correlated activations for noisy images with those for their clean counterparts (Figure 3A, inset), treating the latter as the undistorted reference and reasoning that greater similarity relative to the feedforward backbone reflects effective denoising by recurrence.

**Figure 3:**
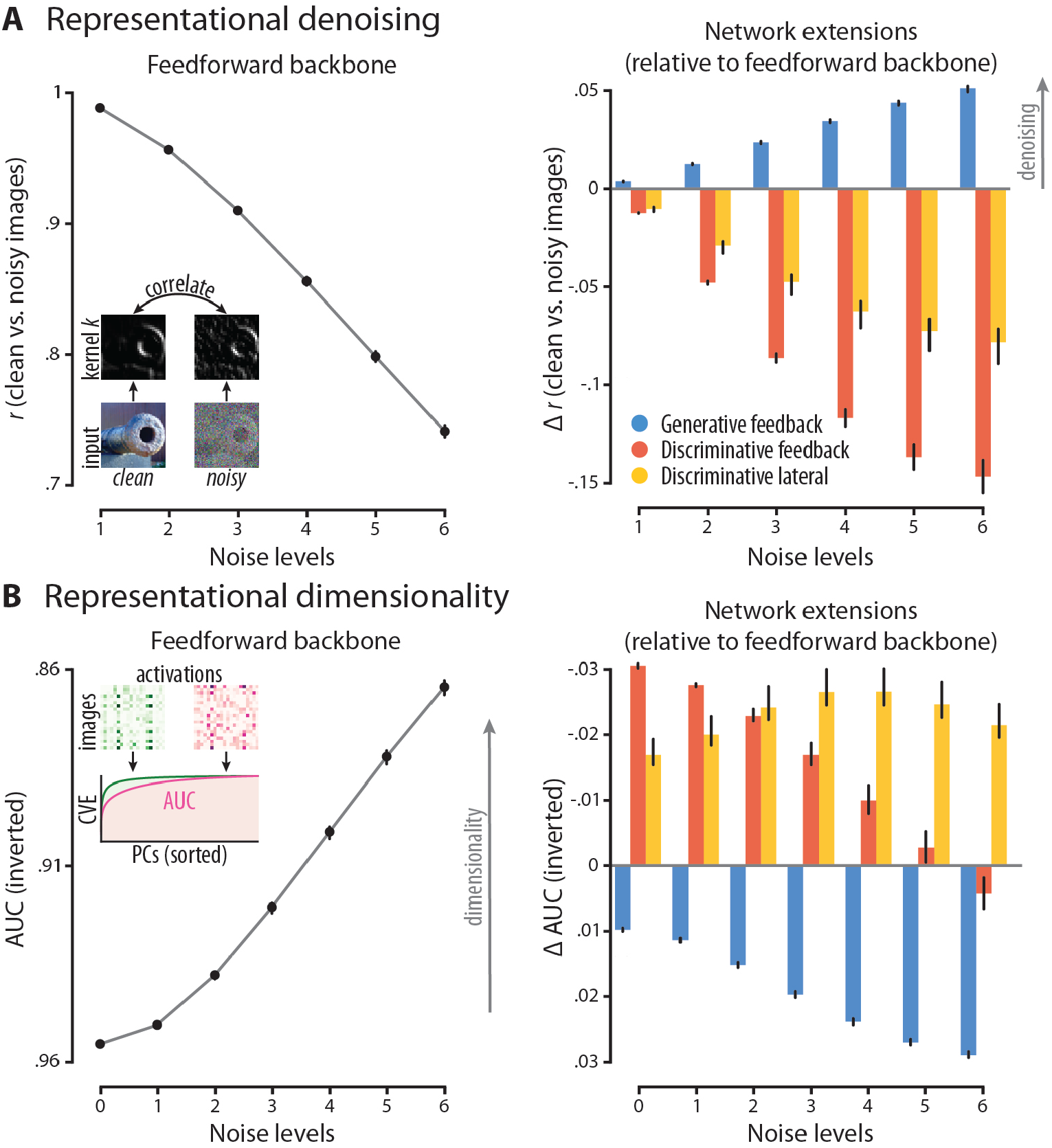
Neural representations at the first encoder. (A) Similarity of activations for noisy images and their clean counterparts (inset), for the feedforward backbone (left) and for recurrent extensions relative to the feedforward baseline (right). (B) Same as (A) for representational dimensionality, measured as the inverted area under the curve (AUC) of the cumulative variance explained (CVE) across principal components (PCs) sorted by explained variance (inset), where the pink example with a smaller AUC than green has higher dimensionality. Error bars represent 95% confidence interval. Generative feedback denoises representations of noisy images and reduces their dimensionality, while discriminative networks do not.

For the feedforward backbone, the noisier an input image, the less its representation resembled that of the corresponding clean image (Figure 3A, left), suggesting that noise substantially changed the feedforward representation of the stimulus. Discriminative networks exacerbated this effect, showing even lower similarity between representations of noisy and clean images than the feedforward backbone, most pronounced for noisier images and the feedback variant. The generative feedback network, by contrast, brought noisy-image representations closer to their clean counterparts than the feedforward backbone, an effect that became most apparent at higher noise levels (Figure 3A, right). This indicates that a generative objective has the capacity to denoise noisy representations, whereas a discriminative objective does not.

### 4.3 Discriminative feedback increases representational dimensionality at low noise, generative feedback decreases it at high noise

We further characterised the geometry of representations by analysing principal components of network activations across images. Representations that are spread across many dimensions require more principal components to account for their variance, and therefore show a smaller area under the cumulative variance explained curve (Figure 3B, inset). As noise introduces additional independent feature dimensions, feedforward representations became more high-dimensional with increasing noise (Figure 3B, left).

On average, a discriminative objective increased representational dimensionality relative to the feedforward backbone, while a generative objective decreased it. This effect was modulated by noise: discriminative feedback raised dimensionality primarily at low noise, discriminative lateral recurrence at intermediate noise, and generative feedback reduced dimensionality most strongly at high noise (Figure 3B, right). The differential consequences of type of recurrent architecture and training objective for representational dimensionality underscore the importance of evaluating representations across the full range of viewing conditions.

### 4.4 Noise training improves classification at intermediate noise for discriminative recurrence

The networks described above were trained on clean images, yet biological vision develops under noisy conditions throughout development. As noise training has specifically been suggested as a biologically plausible method to induce robustness in discriminative networks [Jang et al., 2021], we retrained all network extensions on images degraded at an intermediate noise level and repeated all analyses.

For classification accuracy, recurrent networks showed a smaller overall gain over the feedforward baseline when trained on noisy than on clean images. Discriminative networks showed modest gain for moderately degraded images but lost it for mildly degraded images. The generative feedback network showed no performance gain relative to the feedforward baseline under noise training (Figure 4, right).

**Figure 4:**
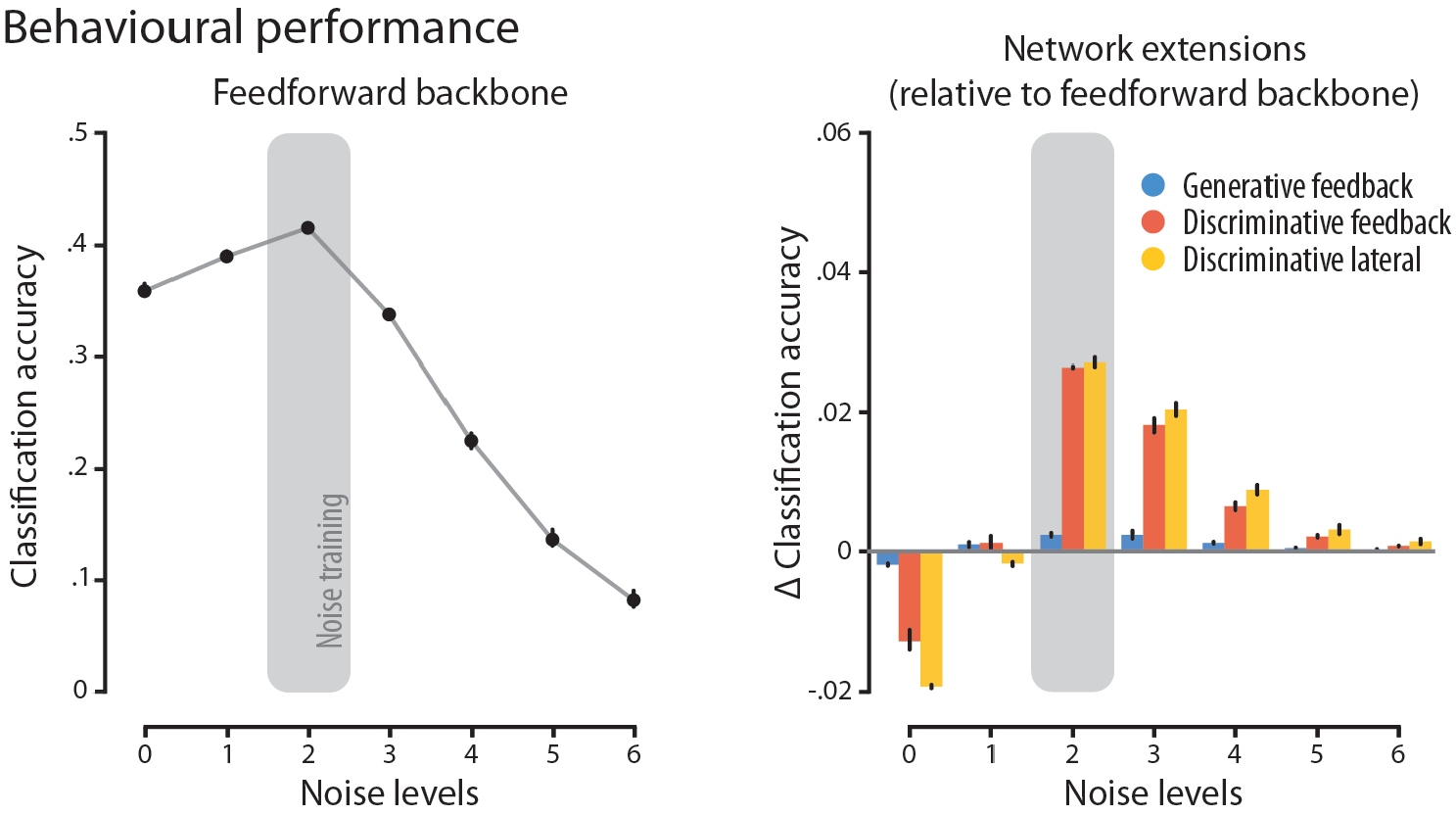
Behavioural performance for noise-trained networks. Classification accuracy of the feedforward backbone trained on images at noise level 2 (grey shading; left). Classification accuracy of recurrent extensions trained on noisy images, relative to the feedforward baseline trained on clean images. Error bars represent 95% confidence interval. For discriminative networks, noise training shifts rather than increases classification performance across noise levels.

For neural representations, the generative feedback network remained largely unaffected by noise training, preserving its dimensionality-reduction and denoising effects. The discriminative networks, however, showed the same qualitative pattern, but substantially reduced in magnitude (Figure 5).

**Figure 5:**
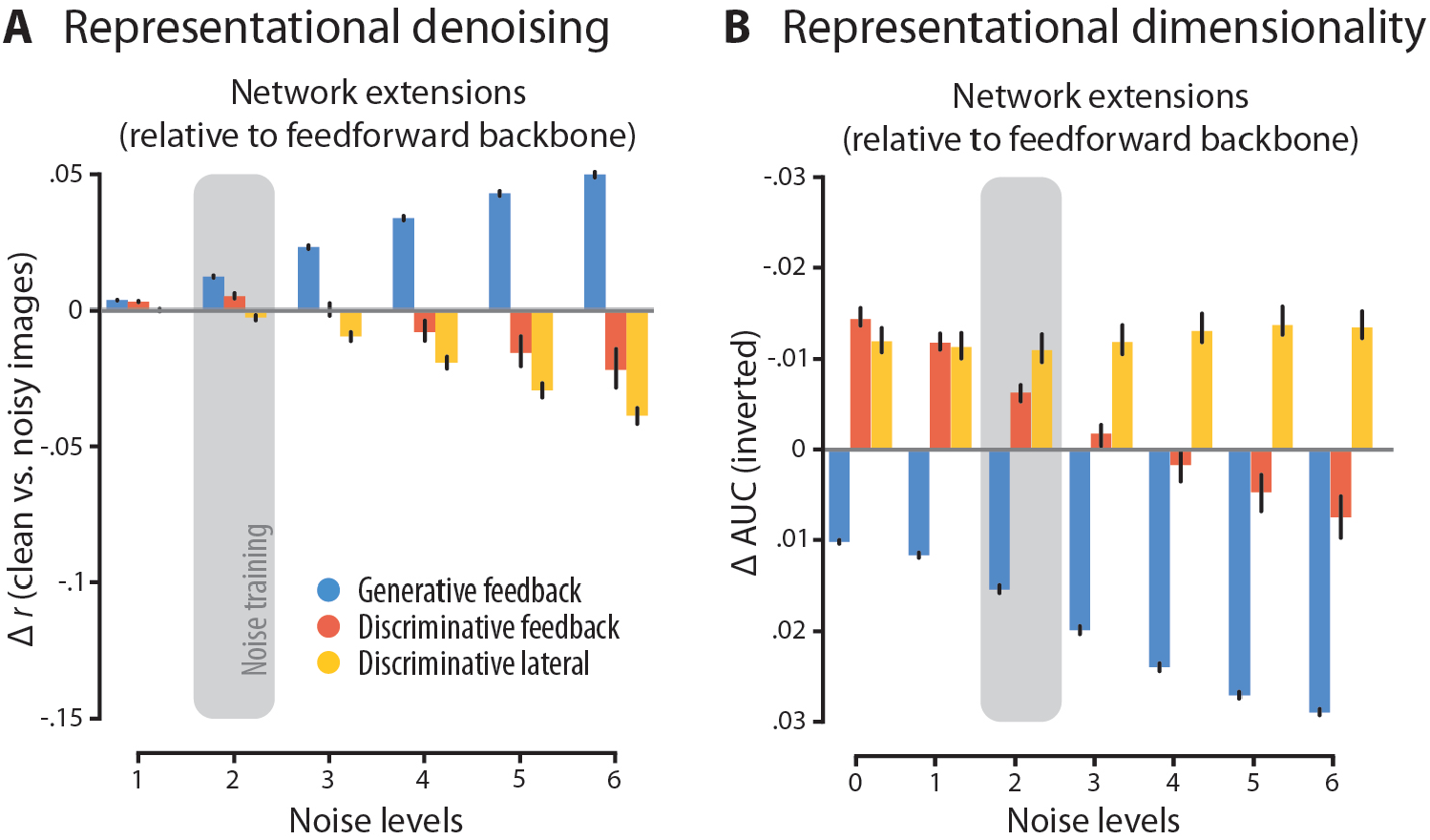
Neural representations for noise-trained network extensions. (A) Representational similarity of recurrent extensions trained on images at noise level 2 (grey shading), relative to the feedforward baseline trained on clean images. (B) Same as (A) for representational dimensionality. Error bars represent 95% confidence interval. Noise training attenuates the representational effects of discriminative but not generative recurrence.

Thus, the performance gain of discriminative networks at the trained noise level came at the expense of performance at other noise levels and their capacity to modulate feedforward representations.

## 5 Discussion

While recurrence is known to improve robustness to noise in biological vision, it is largely unclear how this improvement is achieved. Here, we characterised how different types of recurrent architectures and training objectives reshape noisy feedforward representations. In line with Lindsay et al. [2022], we found generative feedback to reduce dimensionality and improve denoising at the earliest processing stage, whereas discriminative networks with either lateral or feedback recurrence did the opposite — a dissociation that may reflect a fundamental difference in strategy.

Generative feedback embodied a reductionist strategy, where reconstructing low-level from highlevel features may enforce consistency, such that noise components that contradict the broader interpretation are suppressed, yielding more compact representations than feedforward processing as noise grows. By contrast, discriminative networks were expansionist, exploiting features beyond those sufficient for the simple case of clean-image discrimination. Going beyond Lindsay et al. [2022], varying noise levels revealed a further dissociation within discriminative networks. Feedback increased dimensionality most strongly at lower noise levels, consistent with the top-down sharpening of additional features that becomes unreliable as the high-level interpretation itself degrades. Lateral recurrence, by contrast, increased dimensionality most strongly at higher noise levels, suggesting that local integration is particularly apt to recover (or confabulate) additional features when noise is abundant enough to act upon. Though the precise mechanisms remain to be understood, the three recurrent types produce clearly dissociable representational signatures across noise levels that constitute testable predictions for which one the brain employs.

Consistent with theoretical accounts favouring generative objectives for out-of-domain generalisation, discriminative networks performed best at lower noise levels close to the training distribution, while generative feedback generalised to intermediate noise levels more dissimilar from it. This aligns more broadly with the growing success of generative models in machine learning, where the generative objective of diffusion models has been shown to support out-of-distribution image classification [Jaini et al., 2024] and transfer to new video tasks [Wiedemer et al., 2025]. Previous work modelling robustness in the brain found mixed results for generative objectives [Choksi et al., 2020, Lindsay et al., 2022], potentially because MNIST digits are too simple to require the generalisation benefits that become evident under more challenging viewing conditions. We further linked the observed generalisation benefit to a representational signature of strongest denoising at the final encoder, while discriminative networks performed best when dimensionality was highest (see Section A.3). This demonstrates the functional relevance of these representational metrics for robust vision (see Section A.4 for an in-depth discussion).

We rarely encounter ideal viewing conditions, raising the question of whether training on noisy images is a biologically more plausible regime for acquiring robustness. Consistent with noise training benefitting discriminative CNNs [Jang et al., 2021], we observed the same performance benefit and traced it to a boost in the modulatory influence of discriminative recurrence at higher processing stages. That this benefit did not generalise to cleaner images may reflect that noise-trained discriminative networks learn noise-specific rather than noise-invariant object representations. Generative feedback instead preserved its representational modulation under noise training but without behavioural benefit, suggesting that an accurate internal model is best acquired under clean conditions. That discriminative recurrence, requiring exposure to many conditions to learn condition-specific representations, and generative recurrence, requiring high-quality exposure to clean conditions to build condition-general internal models, converge on similar performance at moderate noise levels, broadens the space of testable predictions beyond which type of recurrent architecture and training objective underlie robust vision in the brain to also include under which training regime it is acquired.

### Limitations

First, we used spatially uncorrelated pixel noise, whereas structured noise like blur or occlusion requires more global context and may rely more strongly on generative feedback. Second, we froze feedforward weights, whereas recurrence in the brain may reshape feedforward processing toward features it can act upon more effectively. Third, we isolated individual types of recurrent architectures and training objectives, whereas the brain may employ them jointly (Peters et al., 2024), an interaction outside the scope of the present work.

## Conclusions

We characterised how the type of recurrent architecture and its training objective differentially reshape representations under noise: generative feedback follows a reductionist strategy, achieving behavioural robustness through representational denoising, while discriminative recurrence follows an expansionist strategy, requiring noise training to reach comparable robustness. These dissociable signatures offer a principled framework for interrogating the mechanisms of robust vision in the brain.

## A Technical appendices and supplementary material

### A.1 Supplementary Methods: Hyperparameters

### A.2 Supplementary Results: Behavioural performance and neural representations of

### A.3 Supplementary Results: Representations of the final encoder

The analyses above characterised how recurrence modulates activations at the first encoder. As activations at later encoders become increasingly tuned to the classification decision made at the top of the network [Zeiler and Fergus, 2014], we additionally inspected the final encoder to examine whether the classification gains of discriminative and generative recurrence can be explained by their corresponding modulations of representational denoising and dimensionality.

Compared to the first encoder, denoising in the generative feedback network was now confined to mildly and moderately degraded images, precisely where the classification gain was observed. This was accompanied by a reversal in the direction of the dimensionality effect, with a small increase relative to the feedforward backbone for moderately degraded images. Discriminative networks showed representational effects in the same direction as at the first encoder, but with strongly diminished modulation for severely degraded images, where no classification gain was observed (Figure 6). Together, this suggests that stronger denoising and higher dimensionality relative to the feedforward backbone are the representational signatures of improved classification performance.

**Figure 6:**
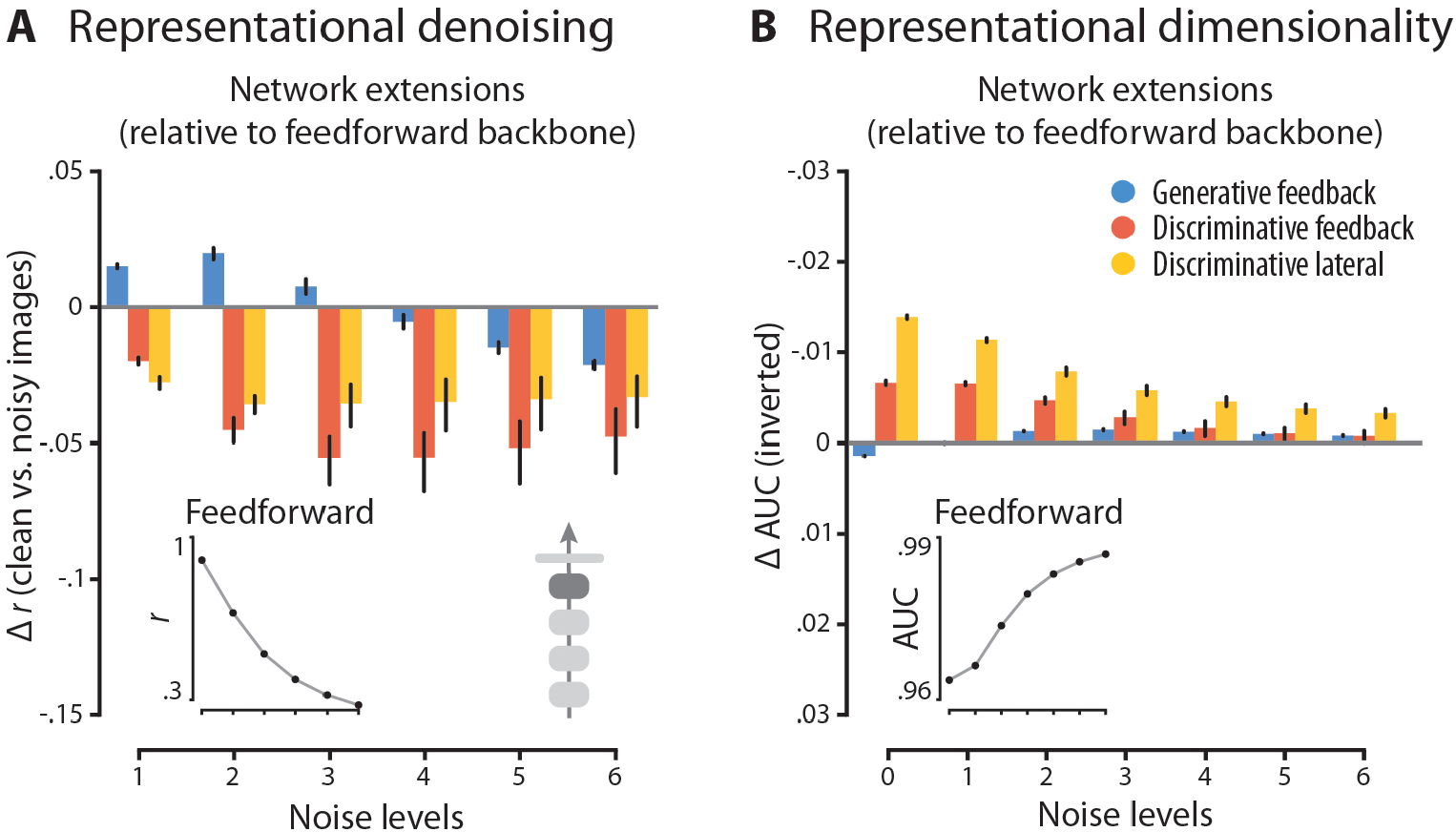
Neural representations at the final encoder. (A) Representational similarity for recurrent extensions at the final encoder relative to the feedforward baseline, across noise levels. Inset shows representational similarity for the feedforward backbone. (B) Same as (A) for representational dimensionality. Error bars represent 95% confidence interval. Denoising and dimensionality increase at the final encoder mark improved classification performance.

### A.4 Supplementary Discussion: Representations of the final encoder

The representational signatures at the final encoder show striking differences from those at the first. Some may reflect pressures intrinsic to the higher processing stage — the absence of dimensionality reduction in generative feedback may reflect pressure to align representations with a more discriminative format, imposed by proximity to the classification decision or by the absence of further feedback. Others may reflect downstream consequences of the early stage — the attenuated denoising for generative feedback and attenuated dimensionality increase for discriminative lateral recurrence at high noise suggest that information added or removed at earlier stages was not of functional relevance at this more abstract level. Taken together, this raises new questions about how recurrent architecture and training objective interact across the processing hierarchy.

